# Evolutionary rates in ‘living fossils’ across the Tree of Life

**DOI:** 10.64898/2026.10.03.756400

**Authors:** Manuela Cascini, E. Karen López-Estrada, David A. Duchêne, Simon Y. W. Ho

## Abstract

Lineages that have undergone little apparent evolutionary change over long geological intervals are often described as ‘living fossils’, with iconic examples including the coelacanth, tuatara, and *Ginkgo*. However, studies of morphological and molecular data have produced mixed evidence regarding whether living fossils have evolved more slowly than their relatives. We tested for reduced evolutionary rates using a sister-pairs approach, analysing a comprehensive set of morphological and molecular data for 45 living-fossil lineages and their close relatives across the Tree of Life. Living-fossil lineages showed substantially reduced rates of morphological evolution, whereas evidence for reduced molecular rates was weaker. To test whether this pattern could be explained by species poverty, we also analysed other depauperate lineages (species-poor clades contrasted with species-rich sister clades) and found a weak tendency towards reduced molecular rates but without a corresponding slowdown in morphological evolution. Our study reveals that the living-fossil concept captures a quantifiable pattern of prolonged morphological conservatism, but also suggests that living fossils lack other shared biological or life-history characteristics that might unite them as a group.

## Main

The concept of ‘living fossils’, evocative of evolutionary stasis, has had enduring appeal in the biological sciences and is familiar even to the general public. Historically placed between metaphor and scientific inquiry, the term appeared in Victorian popular culture with a wide range of biological and non-biological meanings (Lidgard and Kitchen 2025). In *On the Origin of Species*, Darwin (1859) used the term to refer to anomalous freshwater forms such as the platypus and lungfish, suggesting that these have persisted in isolated environments under reduced competition and that they might provide a window into ancient forms of life. Simpson (1949) later described such taxa as ‘immortals’, listing the tuatara, opossum, and *Lingula* brachiopods among his examples. These taxa have since become emblematic of long-term evolutionary persistence and morphological stasis.

In evolutionary biology, living fossils remain a somewhat nebulous concept and have been defined in various ways (Figure 1). The differences between definitions can be substantial: living fossils have been described as taxa that resemble their fossil relatives, have low species richness, occupy ecologically marginal niches, or are phylogenetically isolated. These criteria are often applied in different combinations to classify taxa as living fossils (Grandcolas et al. 2014; Werth & Shear 2014). Thus, it is unclear whether living fossils represent a coherent biological phenomenon or are simply a collection of taxa that sit at one end of the spectrum of evolutionary rates across the Tree of Life (Schopf 1984; Bennett et al. 2017). The most direct evolutionary prediction associated with living fossils is reduced morphological evolutionary rate compared with their close relatives. This expectation has been extended to the genomes of living fossils, with suggestions that they experience slower molecular evolution. Simpson (1944) argued that because phenotypic evolution requires genetic change, rates of phenotypic evolution should be related, although not identical, to rates of genetic modification. However, the evidence for slower molecular evolution in living fossils has been mixed (Rieppel et al. 2023). Some studies have reported low molecular rates in putative living-fossil lineages, including elephant sharks, gars, bowfins, sturgeons, and paddlefishes (Venkatesh et al. 2014; Braasch et al. 2016; Thompson et al. 2021; Brownstein et al. 2024), whereas others have found rates in cycads, tadpole shrimps, and polypterid fishes to be comparable to, or even to exceed, those of their relatives (Nagalingum et al. 2011; Mathers et al. 2013; Near et al. 2014). Coelacanths illustrate this ambiguity: slow evolution has been reported in protein-coding genes (Amemiya et al. 2013), while broader claims of evolutionary stasis in this taxon have been challenged on both morphological and molecular grounds (Casane & Laurenti 2013).

**Figure 1.**
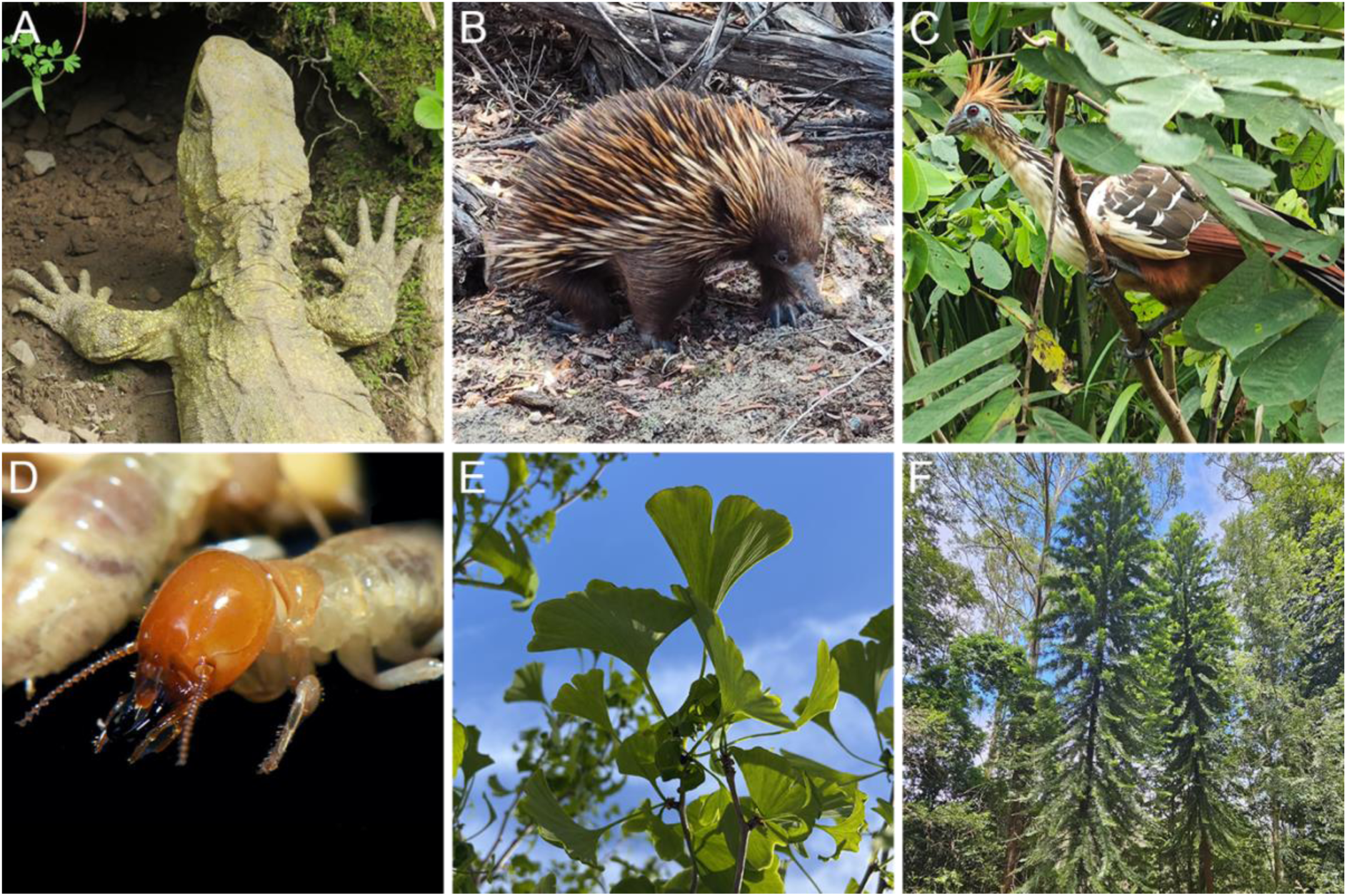
Organisms that have been referred to as ‘living fossils’. (A) Tuatara, *Sphenodon punctatus*. (B) Short-beaked echidna, *Tachyglossus aculeatus*. (C) Hoatzin, *Opisthocomus hoazin*. (D) Giant northern termite, *Mastotermes darwiniensis*. (E) Ginkgo, *Ginkgo biloba*. (F) Wollemi pine, *Wollemia nobilis*. Photographs by S. Ho (A), Z. Zhang (B), Jorge Gutiérrez Rodríguez (C), Y.-K. Tea (D), M. Cascini (E), and R. Dimon (F).

The concept of living fossils is viewed with varying degrees of disapproval among biologists. A common concern is that the term can evoke a ladder-like (*scala naturae*) view of evolutionary history, reminiscent of aspects of Haeckelian thinking, in which living fossils are interpreted as ancient or primitive forms that have somehow stopped evolving (Brouwers 2012; Nuwer 2013; Yong 2013; Carnall 2016). However, some have defended the term as a useful metaphor for integrating different lines of evidence, framing questions about evolutionary stasis, and guiding conservation efforts (Lidgard & Love 2018; Turner 2019; Vargas 2023). There have been elegant attempts to give the concept greater rigour, including through phylogenetic definitions (Turner 2019) and quantitative measures of ‘living fossilness’ (Bennett et al. 2018).

Here we evaluate the living-fossil concept on a broad scale by comparing morphological and molecular rates using a comprehensive collection of data sets that includes 45 lineages described as ‘living fossils’ across the Tree of Life (Supplementary Table S1). We use a phylogenetic approach to compare rates along branches leading to living fossils with those of their close relatives, while accounting for potential sources of bias including the node-density effect and phylogenetic non-independence (Bromham et al. 2002; Venditti et al. 2006; Venditti & Pagel 2008; Hugall & Lee 2007). Because most groups of living fossils display low taxonomic diversity (Stanley 1979), and because some models predict that species-poor lineages evolve more slowly than their more diverse relatives (Bininda-Emonds 2007; Wagner & Estabrook 2014; Bromham 2024; Budd & Mann 2025), we additionally test whether species poverty can account for the evolutionary patterns observed in living fossils.

### Morphological slowdown in living-fossil lineages

Our analyses show that living fossils have experienced lower morphological evolutionary rates than their sister taxa (Figure 2). We detected this pattern by comparing the lengths of the branches leading from the most recent common ancestor to each living fossil and its sister taxon (Supplementary Data Table S3). We quantified the difference as a log-ratio, with values below zero indicating less accumulated morphological change along the living-fossil lineage. To address the potential biases caused by the node-density effect (NDE), whereby branch lengths tend to be underestimated in species-poor lineages (Venditti et al. 2006; Venditti & Pagel 2008; Hugall & Lee 2007), we balanced taxon sampling within each sister-pair comparison and re-estimated branch lengths using the reduced data sets. We hereafter refer to these as the NDE-corrected analyses.

**Figure 2.**
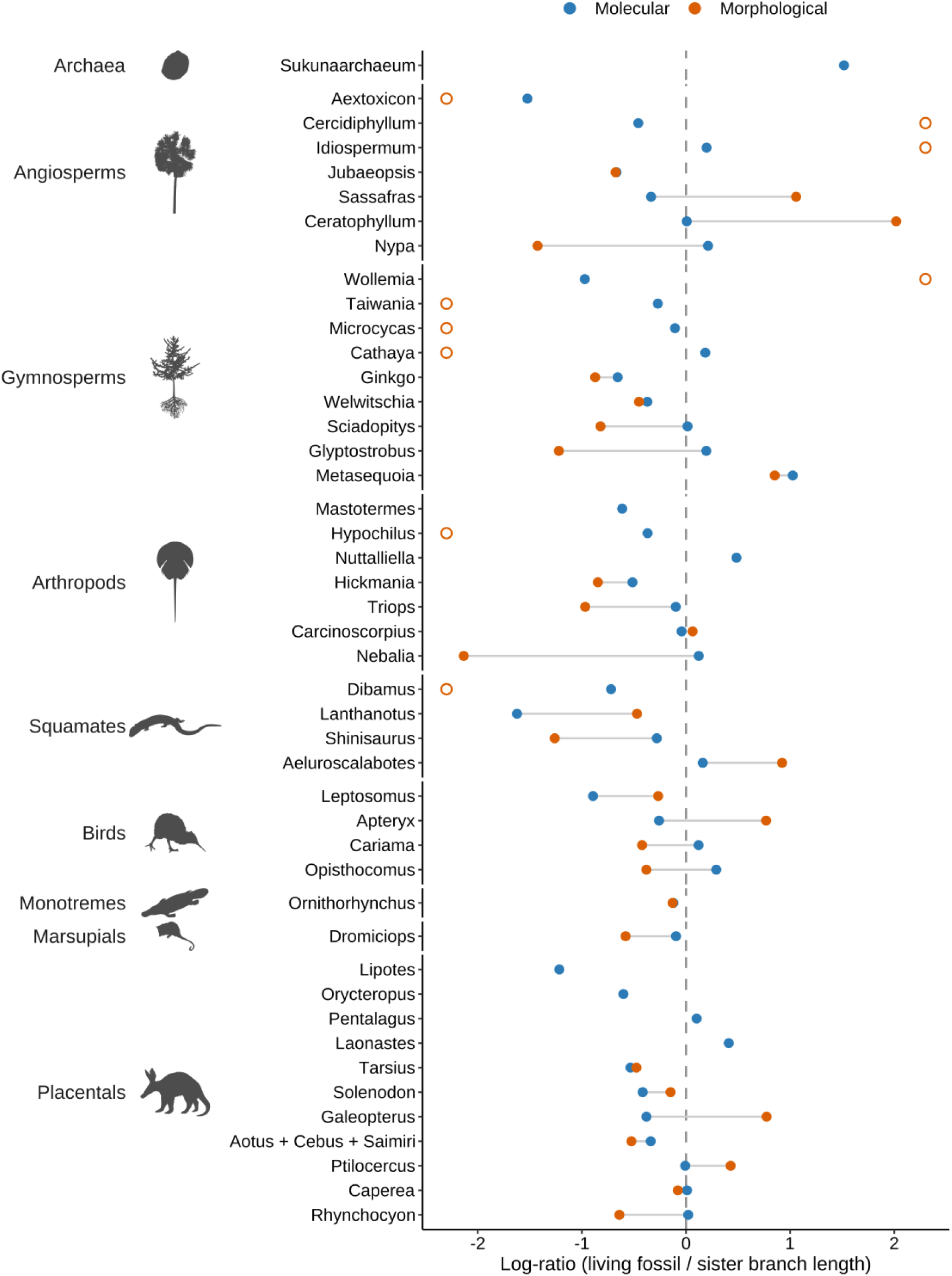
Taxonomic sampling and evolutionary rates of living fossils relative to their sister lineages. Each row is a living-fossil lineage, arranged by taxonomic group to show the breadth of sampling across the Tree of Life. Points show the log-ratio of MRCA-to-tip branch lengths to each living fossil and its sister taxon for morphological data (orange, *n* = 38) and molecular data (blue, *n* = 45), joined by a grey line. Negative values (left of the dashed vertical line) indicate slower evolution along the living-fossil lineage than along its sister lineage. The axis is truncated at ±2.3; nine morphological comparisons fall outside this range and are drawn at the axis limit as open circles, with log-ratios between −13.5 and +13.2. In seven of these, no morphological change could be inferred along one of the two terminal branches, so the magnitude is not interpretable although the direction is. No comparison of molecular change falls outside the range. For *Rhynchocyon*, the morphological value is the joint comparison of *Rhynchocyon* and *Orycteropus* against Paenungulata; the row labelled *Aotus* + *Cebus* + *Saimiri* is a single focal clade in both data types.

In our NDE-corrected analyses, living fossils had shorter morphological branches than their sister taxa in 27 of 38 comparisons (*p* = 0.007, one-sided sign test; *p* = 0.029, one-sided Wilcoxon signed-rank test), with a median log-ratio of −0.47, corresponding to approximately 38% less accumulated change (Figure 3A). The result was robust to removal of any single comparison (*p* = 0.004–0.010, leave-one-out sign tests) and was strengthened when we excluded the seven comparisons in which no change could be inferred along a terminal branch (23 of 31; *p* = 0.005, sign test; *p* = 0.014, Wilcoxon signed-rank test; Supplementary Table S5a).

**Figure 3.**
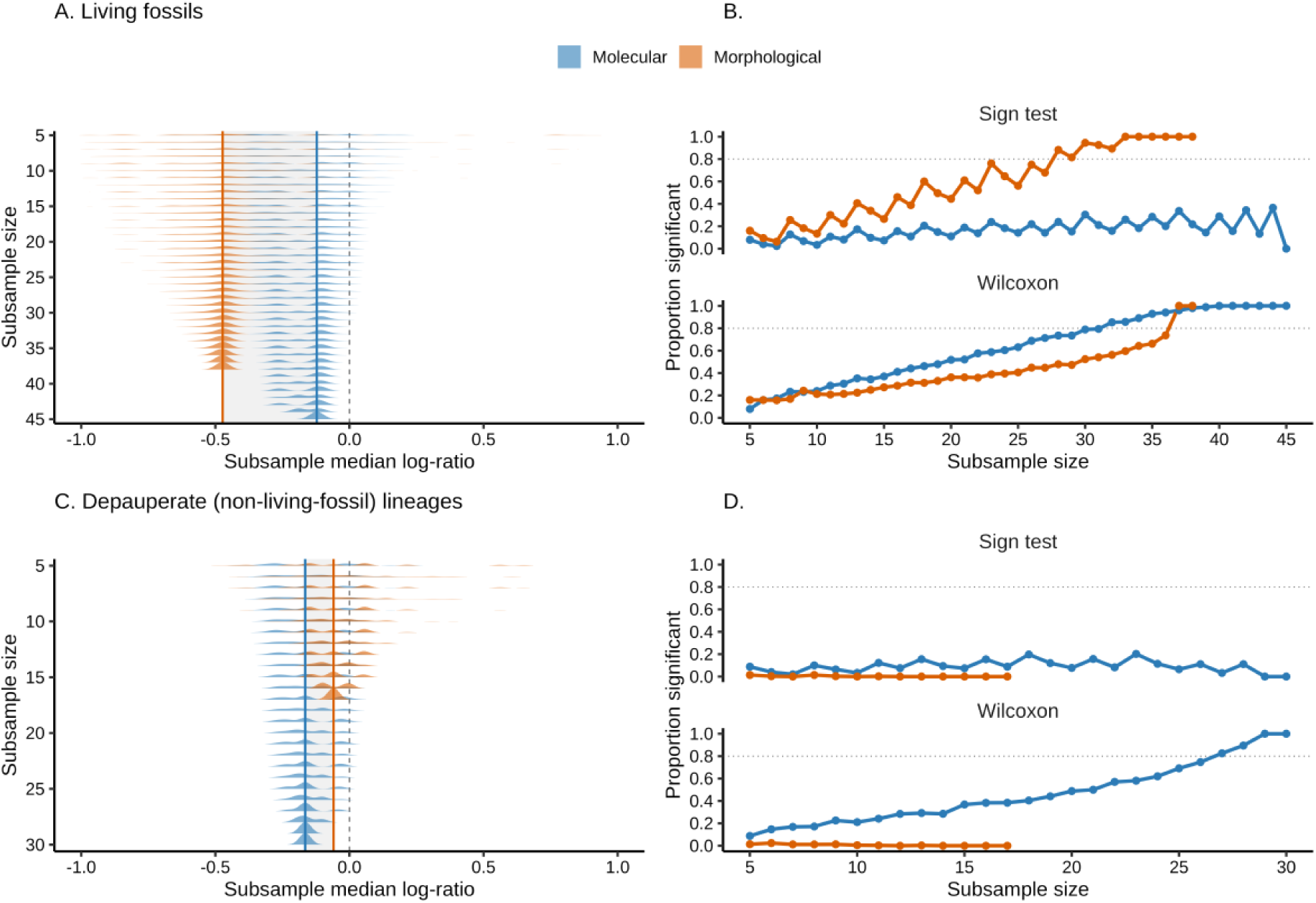
Subsampling robustness of the rate comparisons for living fossils (A, B) and depauperons (C, D). For each lineage, we computed the log-ratio of its MRCA-to-tip branch length to that of its sister representative (or, for depauperon lineages, its more diverse sister representative) under the NDE-corrected analysis; negative values indicate a shorter branch (slower evolution) in the focal lineage. (A) Living fossils. Distributions of subsample-median log-ratios for morphological data (orange; 38 comparisons) and molecular data (blue; 45 comparisons); each ridgeline shows the distribution of 1,000 subsample-median log-ratios drawn without replacement at each subsample size (y-axis). The dashed vertical line marks zero and the solid coloured lines mark the full-data median for each data type. Both distributions lie predominantly below zero, more strongly for morphological data. (B) Proportion of the 1,000 subsample replicates in which living fossils had significantly shorter branches than their sister lineages (one-sided *p* < 0.05) as a function of subsample size, for the sign test (top) and Wilcoxon signed-rank test (bottom). (C) Depauperons. As in (A), for morphological data (orange; 17 comparisons) and molecular data (blue; 30 comparisons); the distribution for morphological data is centred near zero (median −0.06), whereas the distribution for molecular data lies below zero (median −0.16). (D) Proportion of subsample replicates with significantly shorter branches in species-poor lineages (one-sided *p* < 0.05), for the sign test (top) and Wilcoxon signed-rank test (bottom).

**Figure 4.**
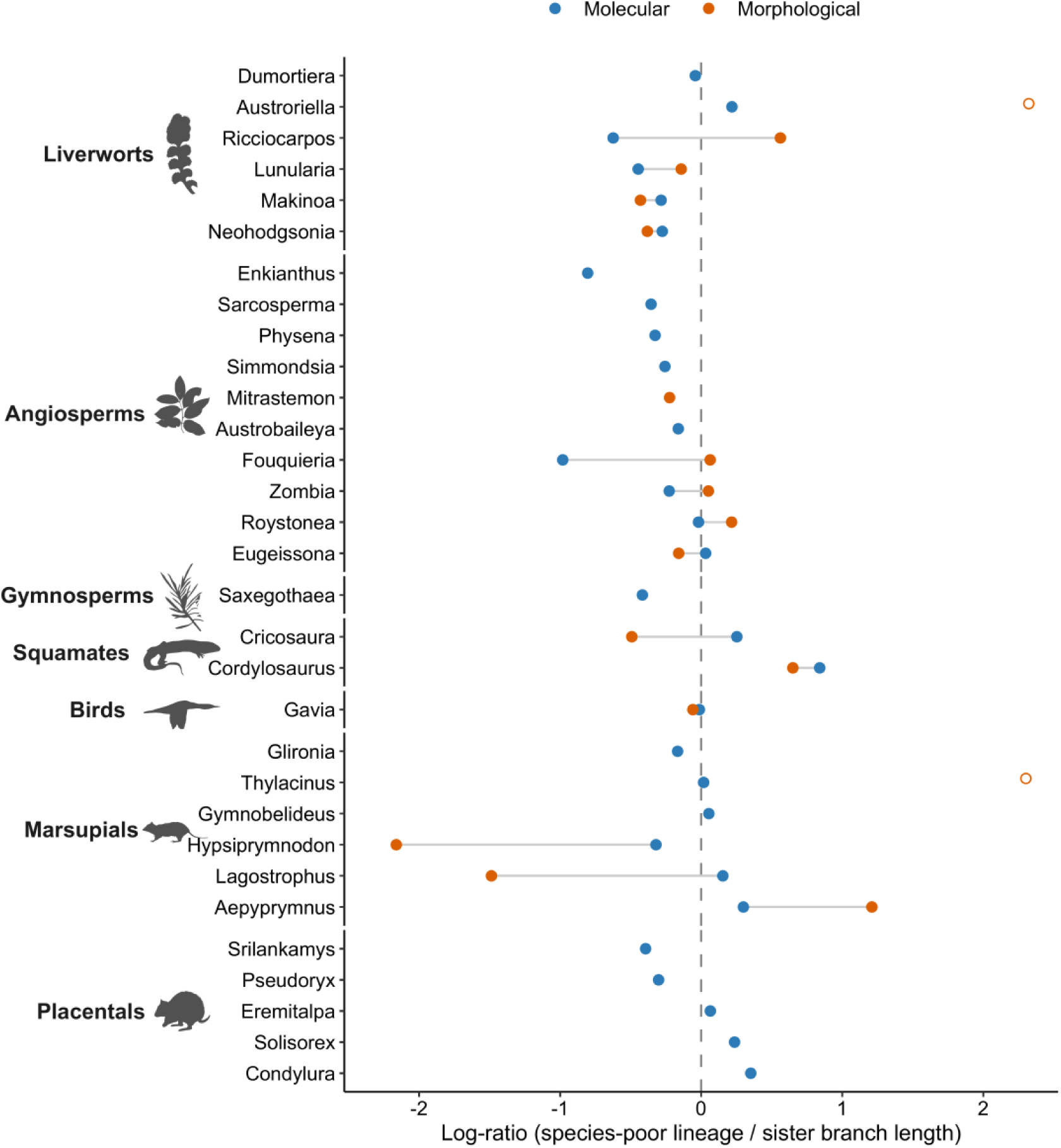
Taxonomic sampling and evolutionary rates of depauperons (species-poor lineages) relative to their sister lineages. Each row is a depauperon lineage arranged by taxonomic group to show the breadth of sampling across the Tree of Life. Points show the log-ratio of MRCA-to-tip branch lengths for the depauperon and its sister taxon for morphological data (orange, *n*= 17) and molecular data (blue, *n* = 30), joined by a grey line. Negative values (left of the dashed vertical line) indicate slower evolution along the depauperon lineage than along its sister lineage. The axis is truncated at ±2.3; two morphological comparisons (*Austroriella*, *Thylacinus*) fall outside this range and are drawn at the axis limit as open circles, with log-ratios between +10.0 and +13.0. In both, no morphological change could be inferred along a terminal branch, so the magnitude is not interpretable although the direction is. No comparison of molecular change falls outside the range.

We focus on the NDE-corrected analyses but compare them with the analyses of the full data sets to assess the effect of uneven taxon sampling. Living fossils also had shorter morphological branches than their sister taxa in our analysis of the full data sets, with 25 of 38 comparisons showing shorter branches (*p* = 0.036, sign test) and a median log-ratio of −0.19 (Figure S1). Thus, the direction of the effect was consistent before and after NDE correction, but with the latter yielding a larger estimated slowdown. The Wilcoxon test provided weaker evidence before NDE correction (*p* = 0.298). Together, these results indicate that morphological slowdown can be detected in the full data set and is not an artefact of the taxon sampling or branch-length re-estimation used to generate the NDE-corrected trees; the correction affects the estimated magnitude of the slowdown but not its direction.

A morphological slowdown in living fossils might appear unsurprising, given that many of these taxa were initially recognised as evolutionary relicts because of their resemblance to fossil relatives (Lidgard & Love 2018). However, only 12 of the 38 taxa in our morphological data set had been designated as living fossils on the basis of morphological conservatism; the remaining 26 taxa were identified as such from relictual distributions (11 taxa), phylogenetic isolation (10 taxa), the rediscovery of a lineage that had been believed to be extinct (3 taxa), or mixed and unclear grounds (2 taxa). Morphological slowdown was evident in both groups: 9 of 12 morphology-based living fossils and 18 of 26 taxa recognised as living fossils on other grounds had shorter branches (median log-ratio −0.47 versus −0.49; *p* = 0.52, Wilcoxon rank-sum test; Supplementary Table S5a). These results suggest that, despite the heterogeneous criteria historically used to identify living fossils, these taxa collectively show a quantifiable reduction in morphological evolutionary rates.

Several mechanisms have been proposed to explain morphological stasis. Genetic or developmental constraints might limit the variation available to selection, thereby restricting the evolutionary trajectories that can be explored by a lineage (Maynard Smith et al. 1985). Nonetheless, morphological conservatism should not necessarily be equated with an absence of phenotypic evolution. Under stabilising selection, phenotypic variation is continually generated but deviations from an adaptive optimum are removed by selection (Charlesworth et al. 1982; Estes & Arnold 2007). Coevolutionary alternation has also been proposed as an ecological mechanism in which changing interactions among species repeatedly alter the direction of selection, potentially producing little net morphological change despite continual evolutionary responses (Kelley et al. 2025).

Morphological change is often mosaic, with different characters and traits experiencing contrasting evolutionary dynamics (Hopkins & Lidgard 2012; Parins-Fukuchi 2020). For example, evolutionary rates can differ substantially among anatomical regions within the same organism, including among cranial regions in birds (Felice & Goswami 2018) and among body regions in ray-finned fishes (Larouche et al. 2018). Conversely, concerted reductions in morphological rate can occur across multiple anatomical elements, as shown by slower evolution across major cranial regions in birds than in non-avian dinosaurs (Felice et al. 2020). An overall reduction in morphological rate can therefore reflect either broad conservatism across the phenotype or a slowdown concentrated in just a few of its parts.

We were able to detect a strong signal of evolutionary slowdown in living-fossil lineages despite pooling morphological characters within each data set. In future studies, partitioning morphological data sets into regions such as cranial versus postcranial characters in vertebrates, or reproductive versus vegetative traits in plants, could determine whether morphological slowdown is concentrated in particular components of the phenotype. Linking such patterns to functional traits, life-history strategies, and other biological characteristics could then provide insight into the precise drivers of morphological slowdown.

### Genomic evolutionary rates in living fossils

Genomic data showed a tendency towards lower evolutionary rates in living fossils, but the evidence was weaker than for morphology. In our NDE-corrected analysis, 28 of 45 sister-pairs showed shorter branches leading to living fossils, with molecular branch lengths being shorter than those of their sister taxa by a median of 11% (*p* = 0.068, one-sided sign test; *p* = 0.005, one-sided Wilcoxon signed-rank test; median log-ratio = −0.12; Figure 3A, B). The signal was stronger in the full data sets, where living fossils had shorter branches in 31 of 45 comparisons (*p* = 0.008, sign test; *p* = 7×10⁻⁴, Wilcoxon signed-rank test; median log-ratio = −0.17), corresponding to 15% shorter branches. Balancing taxon sampling weakened the signal of a molecular slowdown, consistent with a moderate node-density effect but also with some reduction in statistical power.

Morphological and molecular rates showed a weak association, but this relationship was not robust to the choice of statistical test or individual comparisons. Among the 38 living fossils for which we had data to compare morphological and molecular rates, 19 showed reductions in both, eight showed a reduction in morphological rate but not molecular rate, seven showed the reverse, and four showed neither. The log-ratio of molecular branch lengths explained some of the variation in the log-ratio of morphological branch lengths (least-squares regression: *R*² = 0.14, *p* = 0.037; Figure S2). However, this association was not supported by a rank-based correlation (Spearman ρ = +0.07, *p* = 0.66) and was sensitive to individual comparisons: removing any single comparison led to values of *R*² ranging from 0.05 to 0.19, with *p*-values ranging from 0.016 to 0.227.

Early interpretations of stasis invoked genetic homeostasis or genome-wide genetic cohesion, implying that mechanisms maintaining phenotypic stability might also restrict genetic change (Lerner 1954; Mayr 1963; Eldredge & Gould 1972). This assumption underlies a broader expectation of a link between morphological and molecular rates, although recent analyses of eukaryotic data sets have called this into question (Asar et al. in preparation). The mapping between genotype and phenotype is shaped by developmental architecture, pleiotropy, genetic redundancy, and other features of genomic architecture. Substantial molecular divergence can accumulate without much concomitant phenotypic divergence, while relatively few genetic changes can produce large phenotypic differences (Hansen 2006; Wagner & Zhang 2011; Lehner 2013). Stabilising selection acting on plastic traits, for example, can maintain a relatively constant phenotype without necessarily reducing rates of genetic change, and phenotypic accommodation can allow functional phenotypes to persist despite ongoing genetic mutation (West-Eberhard 2005; Watkins 2021). Our results show that morphological conservatism can be seen in living fossils despite the persistence of molecular change.

Living fossils might show a systematic difference in molecular evolutionary rates if they tend to share certain life-history traits that are associated with lower rates, such as long generations, low metabolic rate, longevity, and large effective population size (Bromham et al. 2015; Wang & Obbard 2023). Long generations can involve a smaller number of germline replications per unit time, while effective population size influences the fate of new variants through genetic drift (Lynch 2007). Differences in mutation and DNA-repair processes can also contribute to variation in molecular rates among lineages. These processes provide possible explanations for some of the variation in molecular rates among our comparisons, but testing them directly would require comparable life-history data across extremely diverse taxa with major differences in their reproduction, physiology, or population biology. This diversity also makes it unlikely that any single life-history trait systematically distinguishes living fossils from their sister lineages. For such a trait to explain a general molecular slowdown, it would need to be shared across distantly related lineages of living fossils.

Differences in nucleotide composition provide another possible source of variation in molecular rates. Genomes and genomic regions with lower GC content have been reported to exhibit higher synonymous and nonsynonymous substitution rates than GC-rich groups, potentially reflecting differences in mutation pressures or DNA-repair processes (Lartillot 2013; Bolívar et al. 2019). Across the 43 comparisons based on nucleotide sequences, we found no difference in GC content between living fossils and their sister lineages (median difference = −0.002; *p* = 0.067, Wilcoxon signed-rank test). Furthermore, GC-content differences were unrelated to branch-length ratios (Spearman’s ρ = 0.11, *p* = 0.50).

Although we have found only weak evidence of a broad slowdown in molecular rates along living-fossil lineages, pronounced slowdowns have been documented in individual taxa. Gars have experienced exceptionally low rates of molecular evolution, with highly efficient DNA repair proposed as one contributing mechanism (Brownstein et al. 2024). Similarly, a study of the chambered nautilus (*Nautilus pompilius*) reported low rates of molecular evolution across both coding and non-coding regions of the genome (Zhang et al. 2021). These examples show that extreme molecular conservatism can occur in particular taxa, but our comparative results indicate that it is not a general feature of living fossils.

### Evolutionary rates in species-poor lineages

Living fossils are often species-poor, raising the possibility that their reduced evolutionary rates might be the outcome of some shared driver of lineage diversification and morphological and molecular change. Some models of macroevolution and diversification predict that species-poor lineages accumulate evolutionary change more slowly than their more diverse relatives (Bininda-Emonds 2007; Wagner & Estabrook 2014; Bromham 2024; Budd & Mann 2025). Therefore, we tested whether the signature of an evolutionary slowdown in living fossils is replicated in other species-poor lineages, referred to as depauperons (Donoghue & Sanderson 2015).

We analysed 30 depauperon lineages and found a modest slowdown in molecular evolution but no corresponding slowdown in morphological evolution (Figures 3C, D and 4). For molecular data, 19 of 30 comparisons had shorter branches in the smaller clade, corresponding to a 15% reduction (median log-ratio = −0.165; *p* = 0.100, sign test; *p* = 0.021, Wilcoxon signed-rank test). For morphological data, 9 of 17 comparisons had shorter branches, with a median reduction of 6% (median log-ratio = −0.059; *p* = 0.500, sign test; *p* = 0.555, Wilcoxon signed-rank test). This was much smaller than the 38% reduction in morphological rate observed in living fossils, whereas the difference in molecular rate was similar in magnitude (15% versus 11%). These conclusions were unchanged when we excluded the two morphological comparisons in which no change could be inferred along a terminal branch (9 of 15; *p* = 0.304, sign test; *p* = 0.227, Wilcoxon signed-rank test). Therefore, species poverty can account for some of the pattern seen in molecular rates in living fossils, but does not explain their morphological slowdown.

Depauperons have often been described as more ‘primitive’ than their species-rich relatives. For example, the coelacanth is frequently interpreted and described as an evolutionary relict that retains the morphology of an ancestral lobe-finned fish (Casane & Laurenti 2013). In some extreme cases, terms such as ‘basal’ or ‘ancestral’ are incorrectly applied to extant species-poor lineages. However, such interpretations can be highly misleading, because these lineages have evolved for the same amount of geological time as their more diverse sister lineages since they split from their most recent common ancestor (Krell & Cranston 2004; Beaulieu et al. 2026). Both lineages might display mixtures of plesiomorphic and apomorphic traits, but our analyses fail to find any evidence that depauperons have systematically undergone smaller amounts of morphological or molecular change. Instead, the occurrence of depauperons might simply be an outcome of stochastic differences in diversification, rather than a history of reduced evolutionary change (e.g., Scholl & Wiens 2016; Kopperud et al. 2026). Distinguishing between these possibilities requires comparisons of evolutionary rates rather than inferring evolutionary history from species richness alone.

Whether depauperons are predicted to retain ancestral characters varies across macroevolutionary models. Under simple models without selection, ancestral character states can persist while derived states accumulate elsewhere, potentially increasing the frequency of apparently conservative lineages over time (Raup et al. 1973). Other models instead predict that long-lived, species-poor lineages should be uncommon (Wagner & Estabrook 2014). More generally, relationships between character evolution and diversification might fall on a continuum, with the phylogenetic distributions of character states reflecting different combinations of speciation, extinction, and trait gain and loss (Bromham et al. 2016). Therefore, our comparison tests a specific null expectation: whether the morphological slowdown associated with living fossils is simply a general consequence of species-poor lineage history. The absence of a comparable morphological slowdown in other depauperon lineages argues against this explanation.

### So, what is a living fossil?

The concept of living fossils has been in use for more than 160 years, yet these taxa have been nominated by criteria that are often opaque. The debate over living fossils has been aggravated by a literature built largely on qualitative impressions of individual or few taxa (Sterner 2023). Here, we have moved beyond fragmented and heterogeneous studies by analysing a broad range of cases within a shared quantitative framework. We found a consistent pattern of reduced morphological evolutionary rates in living fossils relative to their sister lineages, a pattern not readily explained by species poverty or other methodological factors examined here. However, this morphological slowdown was not accompanied by a comparable slowdown in molecular evolution. This lack of concordance argues against a general mechanism underlying evolutionary slowdown across living fossils. If living fossils shared life-history traits such as long generations or very low metabolic rates, we would expect to see a stronger effect on molecular rates. Moreover, any such trait would need to recur across the diverse and phylogenetically scattered taxa in our data set, which spans plants, invertebrates, and vertebrates with markedly different biology, ecology, and life history. Instead, our results suggest that living fossils do not constitute a distinct biological category with shared evolutionary properties, but represent the slow end of a continuous distribution of morphological evolutionary rates.

Our findings also have implications for the epistemic value of scientific concepts more generally. As Brigandt & Love (2012) and Brigandt (2020) argue, biological concepts organise knowledge and guide investigative practice, hence their value is not exhausted by their role in classification. Living fossils can therefore be understood as an investigative kind (Sterner 2023): their value lies in directing attention towards unusually low rates of morphological evolution and the processes that produce morphological conservatism, rather than in identifying a biologically distinct category. In this sense, the concept can remain scientifically useful even if there is ultimately nothing biologically special about the taxa that it encompasses.

## Methods

### Taxon sampling

Candidate living fossils were identified through searches of journal articles and scholarly books. We considered any taxon referred to as a living fossil in the scientific literature to be a candidate, regardless of the original rationale for its designation. For each candidate living fossil, we assembled a morphological and/or molecular data set. Because the concept of a living fossil has been applied at different taxonomic levels, any monophyletic group, ranging from species to higher taxonomic ranks, was considered eligible for inclusion following Bennett et al. (2018). We also sampled the sister lineage of each living fossil to enable phylogenetically independent paired comparisons. Accordingly, candidates were required to be lineages with an identifiable sister lineage in at least one data type.

Sister taxa in a pair do not necessarily refer to two sister species. Rather, they refer to two lineages or clades that group together to the exclusion of all other taxa included in the analysis. Where two designated living fossils occurred as sister lineages, they were treated as a single focal clade and compared with their sister clade at their ancestral node; one such case occurred in our analysis of morphological data from mammals, where sengis (*Rhynchocyon*) and aardvark (*Orycteropus*) were placed as sister taxa and were compared jointly against Paenungulata. Likewise, the platyrrhine living fossils *Aotus*, *Cebus* and *Saimiri* form a clade and were compared jointly against the callitrichines in both data types. Our data set included 45 sister-pair comparisons between living fossils and their sister lineages, spanning a broad range of taxonomic groups and evolutionary timescales.

We assembled a second data set consisting of depauperons (species-poor lineages markedly less diverse than their sister lineage, and not themselves living fossils) and their sister lineages. This data set comprised 30 species-poor focal lineages, each paired with a markedly more diverse sister lineage. Species poverty was assessed from described taxonomic richness (Table S2) rather than from phylogenetic sampling: focal lineages contained 1 to ∼17 described species (median 1; e.g. *Enkianthus*, ∼12–17 spp., sister lineage to all other Ericaceae, >4,000 spp.) and were not required to be monotypic, since the criterion is disparity in relative diversity rather than monotypy. We excluded living fossils from this data set.

### Morphological and molecular data sets

Morphological and molecular data were drawn from the compilation of López-Estrada et al. (2025, bioRxiv preprint), which assembled published phylogenetic data sets pairing discrete morphological character matrices with molecular sequence data. We standardised each data set by retaining a single representative per lineage at the focal taxonomic level. Full descriptions of the constituent data sets, their sources, and their preparation are given there, and are summarised for the subset used here in Supplementary Table S4.

We supplemented this compilation with data from four further sources, assembled for the present study to include living-fossil lineages not otherwise represented: the mammal phylogeny of Upham et al. (2019), and phylogenies for termites of the genus *Mastotermes*, ticks of the genus *Nuttalliella* (Mans et al. 2011), and the archaean genus *Sukunaarchaeum* (Harada et al. 2025). These contribute comparisons of molecular rates only. The difference between the 38 morphological and 45 molecular comparisons comprises six lineages with molecular data only (*Laonastes*, *Lipotes* and *Pentalagus* from Upham et al. 2019, *Mastotermes*, Nuttalliella and *Sukunaarchaeum*) and the separate molecular comparisons for *Rhynchocyon* and *Orycteropus*, which form a single clade in the morphological analysis. In total, 13 data sets contributed at least one sister-pair comparison each.

### Estimation of branch lengths

We used a phylogenetic sister-pairs approach to study the difference in evolutionary rate between the branches leading to living fossils and to their sister taxa. By definition, any pair of extant sister taxa have the same age (Eldredge 1984), such that any difference in branch lengths (amount of evolutionary change since their most recent common ancestor) reflects difference in the evolutionary rate. A lower rate of evolution in the living-fossil lineage is indicated by a shorter branch compared with that leading to its sister taxon.

We estimated maximum-likelihood branch lengths using IQ-TREE, retaining the data-partitioning schemes, evolutionary models, and topological constraints of López-Estrada et al. (2025, bioRxiv preprint). To minimise potential bias from the node-density effect (Hugall & Lee 2007; Venditti et al. 2006; Venditti & Pagel 2008), whereby branch lengths can be underestimated in sparsely sampled clades relative to densely sampled ones, we pruned each data set to retain only the living fossil lineage, a single representative of the sister clade, and the outgroup taxa present in the original data set. When the sister clade contained multiple taxa, the representative was selected as the taxon whose median patristic (cophenetic) distance to the focal lineage was closest to the median among all sister taxa. Representatives were chosen from distances on the complete tree, but all branch lengths used in downstream comparisons were re-estimated after pruning. For comparison, we also performed analyses on the full trees.

### Statistical support

For each sister-pair of taxa, the inferred branch lengths leading to their most recent common ancestor (MRCA) were compared between the living fossil and its sister taxon. We computed this as the log ratio of branch lengths, so that negative values indicate a shorter branch leading to the living fossil.

Two one-tailed tests were used to assess whether living-fossil lineages had lower evolutionary rates than their sister clades. The sign test scores each comparison by the direction of the difference alone, ignoring magnitude, with the one-tailed alternative testing whether negative differences occur more frequently than positive differences. The Wilcoxon signed-rank test additionally incorporates the magnitude of the log-ratios, testing whether they are lower than zero. We treat the sign test as our primary analysis because it is insensitive to the precision of the underlying branch-length estimates, and the Wilcoxon signed-rank test as a secondary, magnitude-based test. Comparisons between independent groups of lineages used two-sided Wilcoxon rank-sum tests. Robustness was assessed by leave-one-out analysis, with the sign test being recalculated with each comparison removed in turn.

No comparison was excluded from the analyses. In seven comparisons based on morphological data (NDE-corrected analyses; three in the full data sets), however, no change could be inferred along one of the two terminal branches: its length was estimated at or below 10⁻⁴ changes per character, effectively the smallest value that can be returned by the optimiser. The log-ratios of these comparisons should not be interpreted as magnitudes, since their size depends on where that bound lies rather than on the data; they are identified in Supplementary Table S5b together with the character data behind each. Excluding them strengthens the results of both tests (Supplementary Table S5a). No comparison of molecular data is affected, because we did not obtain any zero-length molecular branches.

We performed a subsampling analysis to evaluate the robustness of the inferred rate differences to sample size. For each subsample size, from five comparisons up to the full data set, we drew 1,000 random subsamples of comparisons without replacement and recalculated the sign test and one-sided Wilcoxon signed-rank test for each, recording the median log-ratio. For each subsample size, we then calculated the proportion of subsamples in which living-fossil lineages had significantly shorter branches than their sister taxa (*p* < 0.05). A consistent signal is expected to yield an increasing proportion of significant subsamples as sample size increases, whereas an unstable or highly variable pattern would not. Analyses were performed separately for the full and NDE-corrected data sets to evaluate the impact of sampling bias.

GC content was calculated for each taxon from the nucleotide alignments as the proportion of unambiguous nucleotides that were G or C, and summarised for each comparison as the median across the focal taxa and across the sister representatives, matching the pipeline that we used to analyse branch lengths. We excluded two data sets that comprised amino-acid alignments, leaving 43 of the 45 comparisons. Differences in GC content between paired lineages were tested against zero using a Wilcoxon signed-rank test, and their relationship with the branch-length log-ratio was assessed using Spearman rank correlation.

## Supplementary Figures

**Figure S1.**
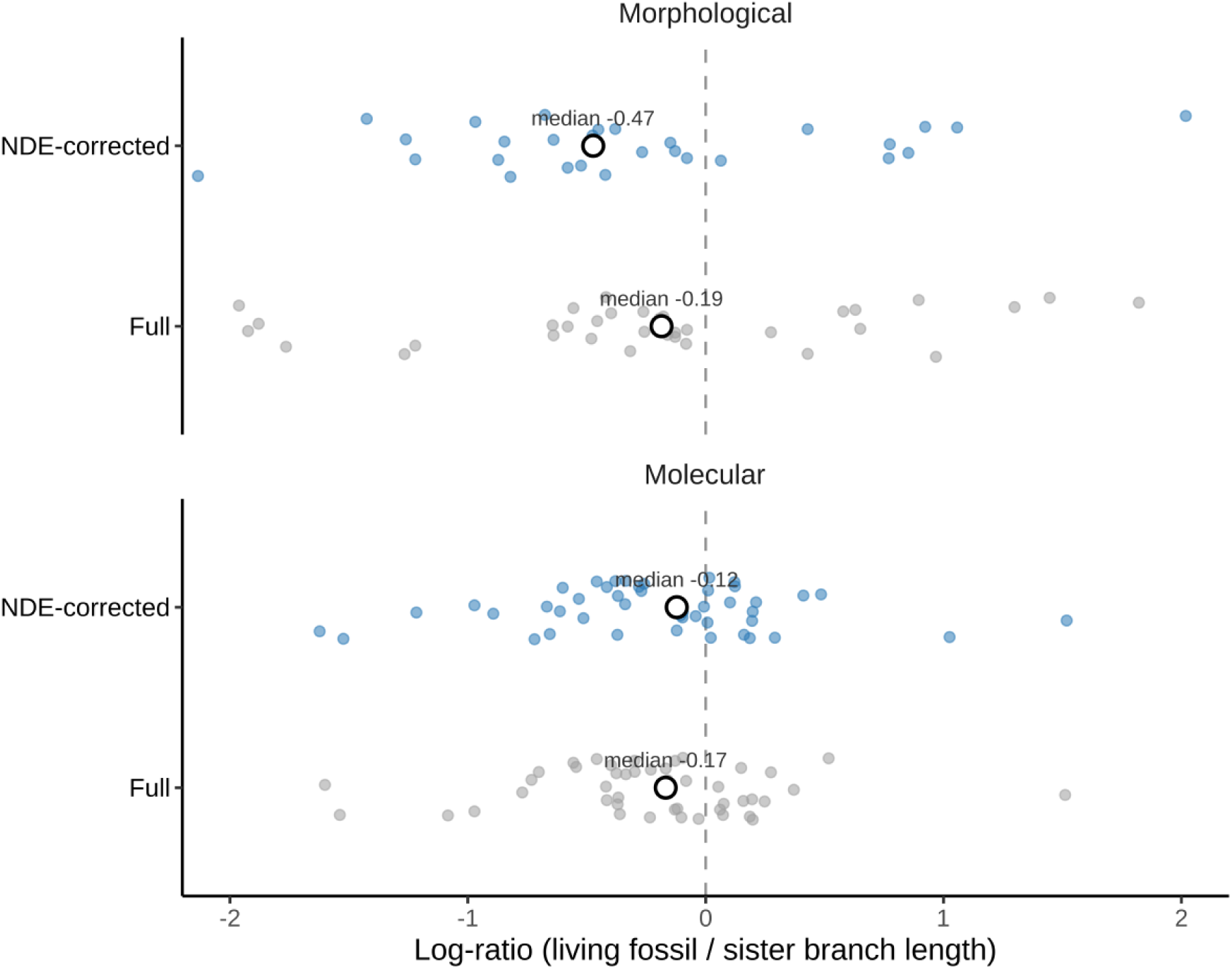
Effect of the node-density correction on the comparison of branch lengths leading to living fossils and their sister taxa. Distributions of the log-ratio of branch lengths (living fossil ÷ sister lineage) for morphological (top) and molecular (bottom) data, from the analysis of the full data (uncorrected) and after the node-density correction.

**Figure S2.**
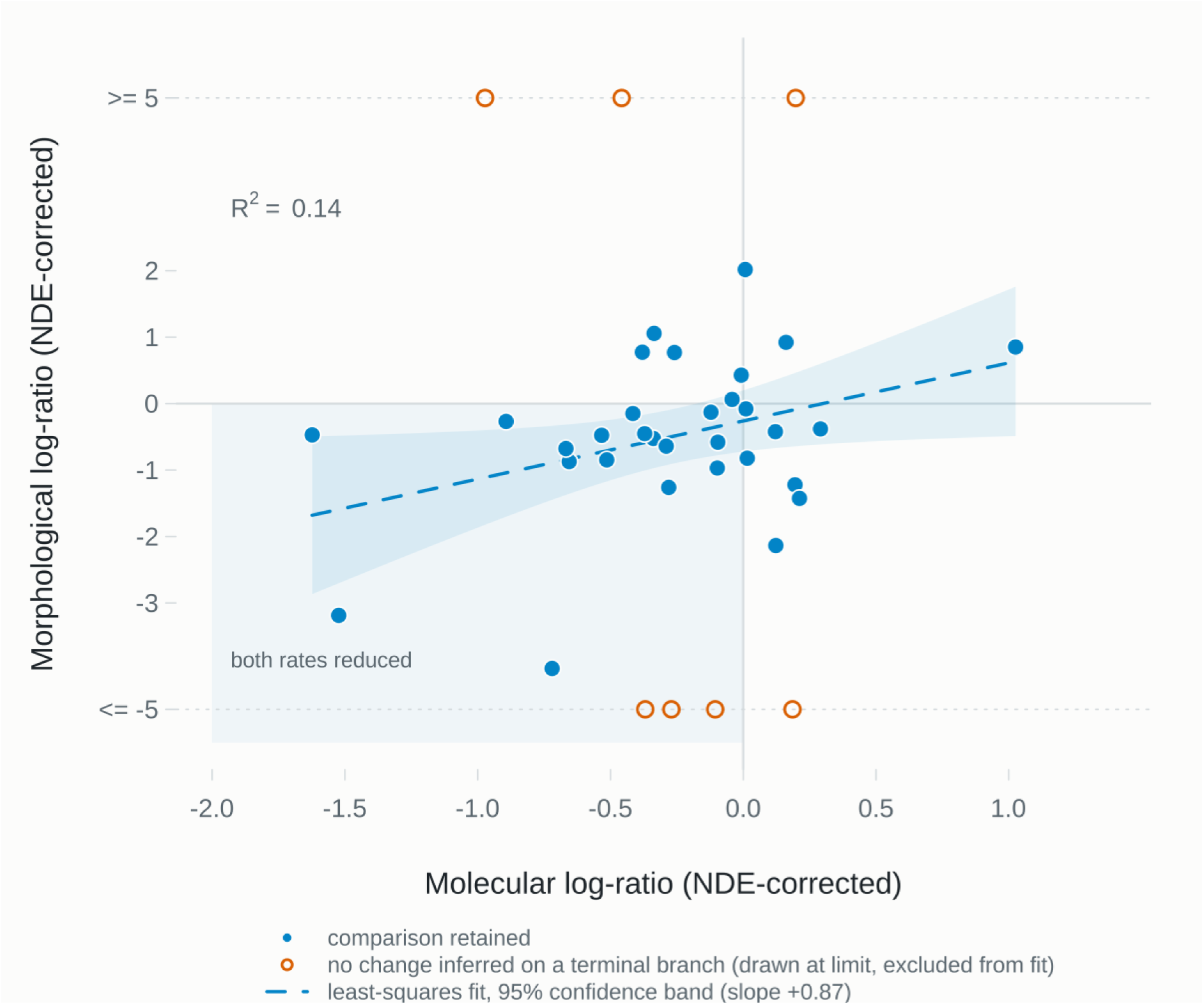
Reductions in morphological and molecular rates are only weakly related across living-fossil lineages. Each point represents one of the 38 living fossils for which we could estimate log-ratios of NDE-corrected morphological and molecular branch lengths. Negative values indicate less accumulated change in the living fossil than in its sister representative, so the shaded quadrant contains lineages that have reductions in both rates. Open circles represent the seven comparisons in which no change could be inferred along a terminal branch. Dashed line and shaded band show the least-squares fit with 95% confidence interval, computed on the 31 finite comparisons only (slope +0.87, *R*² = 0.14, *p* = 0.037; leave-one-out *R*² 0.05–0.19, *p* = 0.016–0.227). Spearman correlations were *ρ = +0.07, p = 0.66* across all 38; *ρ = +0.21, p = 0.25* across the 31 finite comparisons.

